# Negligible participation in the scientific literature of AI unicorn startups

**DOI:** 10.64898/2026.07.15.738744

**Authors:** Quentin E.A. Loisel, Alejandro Sandoval-Lentisco, John P.A. Ioannidis

## Abstract

Artificial intelligence (AI) development is concentrated within private firms, yet their participation in scientific publishing remains poorly understood. We conducted a bibliometric analysis of all 317 AI unicorn startups (1998–2025). Only 1,389 eligible peer-reviewed publications and 688 preprints involving some leading startup contributions were identified. More than half of startups (52.4%) produced no qualifying scientific output, and only 24 firms (7.6%) produced any highly cited papers (≥200 citations). Scientific influence was highly concentrated: the top 10% of firms accounted for 96.8% of citations, while three startups accounted for 92 of 134 firm-attributed highly cited papers. Firm valuation was not associated with publication productivity or highly cited output, whereas funding raised showed weak associations. Overall, participation in formal scientific communication among AI unicorn startups is negligible, comprising only 0.1% of the overall AI literature in 2025. Most leading developers of AI technology do not engage with the scientific literature, raising concerns for the transparency, reproducibility, and accountability of this rapidly moving innovation frontier.

## Introduction

Artificial intelligence (AI) research has increasingly shifted from universities and public laboratories toward private firms with access to large-scale computing infrastructure, proprietary data, and concentrated technical talent (*1*). Commercial AI laboratories and startups now play a central role in developing frontier models and capabilities that influence scientific research, software engineering, medicine, education, and public communication (*2*). In parallel, industry participation in large-scale model development and benchmark performance has increased substantially over the past decade (*1, 3*).

Scientific publishing remains a central mechanism through which research findings are disseminated, scrutinized, and incorporated into cumulative bodies of knowledge. Publications establish scientific priority, facilitate independent evaluation, support reproducibility, and provide a public record of technical advances. As frontier AI development becomes increasingly concentrated within private organizations, understanding whether these firms participate in formal scientific communication has implications for the visibility, transparency, and collective development of knowledge.

Publication activity may also serve functions beyond scientific dissemination, including recruitment, signaling technical leadership, ecosystem influence, and strategic positioning in rapidly evolving competitive markets. Concerns regarding transparency, reproducibility, and governance have become increasingly prominent as frontier AI systems become more capable and socially consequential [3]. Similar concerns have been raised in other innovation-intensive sectors, where highly valued startups, such as Theranos (a case of fraud) and many apparently legitimate unicorns, have sometimes claimed to generate substantial technological advances while producing little or no peer-reviewed scientific evidence (*4*–*6*). Despite growing interest in the capabilities, valuation, and governance of frontier AI firms (*3, 7*), their participation in the scientific literature remains poorly characterized.

Here, we systematically examined the scientific publishing activity of AI unicorn startups, defined as privately held AI companies valued at US$1 billion or more. Using publication and preprint records retrieved from Clarivate databases, we identified scientific outputs related to the AI field and associated with 317 AI unicorn startups. To distinguish substantive startup participation from broader collaborative involvement, firm-level analyses focused on publications with leading authorship contributions, defined as first- or last-author status, or alphabetical order when author order. We evaluated participation in scientific publishing, scientific productivity, citation impact, production of highly cited papers (≥200 citations), publication and citation concentration, author-affiliation patterns, and the relationships between scientific publishing activity, financial scale, and geography.

## Materials and Methods

### Study design and data sources

We conducted a cross-sectional bibliometric analysis of scientific publishing among artificial intelligence (AI) unicorn startups. The study population comprised AI startups valued at ≥US$1 billion identified from two public unicorn lists: Failory.com “The Full List of 308 Artificial Intelligence Unicorn Startups (2026)” (last updated 18 December 2025; accessed 04 February 2026) and Eqvista “Top AI Startups by Valuation” (published 11 November 2025; accessed 04 February 2026). The lists were merged and deduplicated before publication retrieval. The publication window covered 1998–2025, corresponding to the founding year of the oldest company in the cohort through the last complete study year.

Before retrieval, startup names were manually reviewed to identify prior company names, rebranding events, and acquisitions that could affect affiliation matching. This process identified 17 historical name variants corresponding to 14 startups and revealed one duplicated company listed under an alias, reducing the final cohort from 318 startups in the protocol to 317 analyzed firms.

Bibliometric records were retrieved programmatically through the Clarivate Web of Science Expanded API across the Web of Science Core Collection, INSPEC, and the Preprint Citation Index. Searches were conducted using startup names and identified variants in the Address field (AD=“{startup name}” AND PY=(1998-2025)). Address-field retrieval was used because many startups were absent from the databases’ institutional repertoires.

One startup (“Kong”) was excluded from automated retrieval because its query generated excessive nonspecific matches that disrupted API execution. Manual searches of the databases and the company website identified no eligible publications.

Retrieved records included the Web of Science identifier, DOI, title, publication year, authors, affiliations, abstract, keywords, document type, and citation counts. Citation counts corresponded to the platform-wide “All Databases” metric when available. Duplicate records arising from multiple startup variants or overlapping databases were resolved hierarchically using normalized DOI, normalized title, and Web of Science identifiers while preserving all startup and database matches.

## Data preparation

After downloading, the Web of Science Core Collection (n = 116,233), INSPEC (n = 22,619), and the Preprint Citation Index (n = 5,333) datasets were merged, yielding 144,185 records. Five filters were applied sequentially to prepare the data for analysis.

First, a Boolean keyword filter was applied to titles and abstracts to retain records related to artificial intelligence. The comprehensive search string used to identify broader trends in artificial intelligence research is reported in Supplementary Material 3. This filter excluded 129,372 records.

Second, records were restricted to eligible document types: articles, proceedings papers, reviews, journal papers, conference papers, conference proceedings, and preprints, with database-specific term variations harmonized. Retracted records and ineligible document types were excluded. This step removed 302 additional records.

Third, records were excluded if they were published after 2025 or before the matched startup’s founding year. This step removed 1,795 additional records, including 80 records dated after 2025 and 1,715 records predating the relevant startup founding year. These exclusions were highly concentrated among startups with generic or ambiguous names where other entities may have had a shared name: 10 startup names accounted for 89.5% of all date-based exclusions, representing 1,606 of 1,795 excluded records. The largest source of ambiguity was “Cognition,” which accounted for 43% of these exclusions because the term is commonly used in cognitive-science institute names, such as “Centre de Recherche Cerveau et Cognition,” “Institute for Cognition and Information,” and “Institute for Cognition Research and Information

Technology.” “Harvey” accounted for 12% of exclusions and primarily reflected matches to surnames or institutional fragments, such as “Harvey Mudd College, Department of Engineering.” “Sierra” also accounted for 12% of exclusions and frequently matched place names or unrelated organizations sharing the term, including “University of Arizona, Sierra Vista” and “Sierra Nevada Corporation.” Other recurrent sources of false-positive matches included “Ada” and “Turing” (5% each), “Gamma” and “Plus” (4% each), “Pilot” (3%), “xAI” (2%), and “Eve” (1%), which overlapped with place names, personal names, generic organizational terms, technical terminology, or unrelated research-center names.

Fourth, duplicate records were removed using database priority rules and bibliographic similarity. Web of Science records were prioritized over INSPEC records, which were prioritized over Preprint Citation Index records. Duplicates were identified using DOI matches, exact normalized title matches, and fuzzy title/abstract similarity; fuzzy duplicates required title and abstract similarity of at least 90%, or title similarity of at least 95% when abstracts were unavailable. This step excluded 3,248 additional records.

Finally, startup affiliation matching was performed using a curated affiliation dictionary mapping institutional variants to canonical startup names. Institution names were extracted from affiliation/address strings, converted to lowercase, and matched exactly to dictionary entries; prefix and substring matching were not used to reduce false positives from generic or overlapping names. This final filter excluded 6,058 additional records.

The five filtering steps yielded 3,410 records, which have been included for subsequent startup-affiliation identification.

### Attribution of authorship and leading role in authorship

To identify the contribution and the potentially leading role of AI unicorn startups in publications, author order and affiliation data were extracted from the 3,410 records that passed the five cleaning filters. First, papers were classified by whether authors were listed alphabetically, as alphabetical ordering can obscure conventional assumptions about author contributions and leadership. Of the included papers, 3,153 (92.5%) were not alphabetically ordered, 195 (5.7%) followed alphabetical author ordering, and 55 (1.6%) were single-author publications for which alphabetical ordering was not applicable. A further 7 records (0.2%) could not be classified for author ordering because author-position metadata was not parseable.

Second, startup affiliations were identified for authors occupying first, middle, and last author positions. Three variables were created to indicate whether a startup-affiliated researcher occupied each position and, when applicable, to record the corresponding startup. Overall, 1,559 papers (45.7%) included a startup-affiliated first author, 2,808 (82.3%) included at least one startup-affiliated middle author, and 1,617 (47.4%) included a startup-affiliated last author. A total of 1,135 papers (33.3%) contained both a startup-affiliated first author and a startup-affiliated last author.

Third, a lead-startup attribution procedure was developed to assign publications to a startup when multiple startup affiliations were present. For each paper, the number of startup-affiliated authors associated with each startup was counted, and the publication was attributed to the startup represented by the largest number of affiliated authors. Among the 1,135 papers with both startup-affiliated first and last authors, 1,128 (99.4%) had the same startup represented in both positions and therefore presented no first-last attribution conflict. Seven papers (0.6%) involved different startups in the first- and last-author positions. Of these, five were resolved to a single lead startup based on the number of affiliated authors, whereas two remained tied. Across the full corpus, 24 publications (0.7%) had an equal number of affiliated authors across competing startups and were classified as “EVEN”. For these cases, the attribution was shared: each tied startup received one paper and the associated citations in firm-level analyses, while the publication itself was counted only once in corpus-level totals. Five records were found with corrupted CSV fields and so were unrecoverable.

Finally, startup aliases, mergers, acquisitions, and name changes were harmonized using a predefined crosswalk of alternative company names. Publications identified under historical or alternative startup names were reassigned to their corresponding canonical company names to avoid double counting and ensure consistency across analyses. In total, 44 publications (1.3%) were reassigned through this process, all corresponding to records initially identified under Grammarly and subsequently consolidated under Superhuman.

### Records eligibility

Firm-level analyses focused on publications demonstrating substantive startup involvement with a leading contribution to the publication. Accordingly, records were included when they contained at least one affiliation matching a target AI unicorn startup and met one of the following criteria: (1) a startup-affiliated researcher occupied an authorship position suggesting a leading contribution, defined as first or last author (n=2,041); (2) the publication used alphabetical author ordering, making positional authorship less informative (n=195); or (3) the publication was classified as an EVEN case (n=24), in which multiple startups had identical numbers of affiliated authors and no unique lead startup could be assigned.

Of the 3,410 records that passed the data-cleaning and affiliation-validation procedures, 2,077 (60.9%) met the eligibility criteria and were included in firm-level analyses. The remaining 1,333 records (39.1%) contained startup-affiliated authors exclusively in middle-author positions and were classified as collaborative-only publications. These records were excluded from firm-level analyses but retained for analyses examining broader startup participation and collaboration patterns within the scientific literature.

### Author-level affiliation characteristics

To characterize the institutional profiles of startup-affiliated researchers and assess the extent to which scientific contributions were produced by authors working exclusively within startups or through cross-sector affiliations, author-level affiliation characteristics were derived from the *Authors & Affiliations* field of each publication. Affiliation strings were parsed into individual authors and institutional affiliation segments. Institution names were extracted as the text preceding the first comma in each affiliation segment and matched, after standardization to lowercase, against the curated dictionary of AI unicorn startup affiliations.

Each author associated with at least one startup affiliation was retained and aggregated into a unique author-level record. For each author, the startup affiliation(s), observed institutional affiliations across all publications, and authorship positions were recorded. Authorship positions included first author, middle author, last author, and authorship on alphabetically ordered papers. Lead authorship was defined as occupying either the first- or last-author position.

Authors were classified into three affiliation profiles based on all observed affiliations across the corpus. Authors whose affiliations were exclusively associated with startup organizations were classified as “*startup-only”*. Authors holding both startup and academic or research-institution affiliations were classified as “*startup+academia”*. Authors with startup affiliations and other non-academic, non-startup affiliations were classified as “*startup+others”*. When both academic and non-academic affiliations were observed, authors were assigned to the “*startup+academia”* category.

### Outcomes

The primary outcome was the proportion of AI unicorn startups producing at least one eligible highly cited publication. Highly cited publications were defined a priori as eligible publications receiving at least 200 cumulative citations at the time of data collection. Eligible publications were restricted to records involving startup authorship suggesting a leading contribution (first author, last author, or alphabetical authorship where applicable) or shared startup attribution in EVEN cases. A sensitivity analysis excluding alphabetically ordered author lists was conducted.

Secondary outcomes characterized the scale, influence, and patterns of scientific publishing among AI unicorn startups. Measures of scientific productivity included the total number of eligible publications per startup, the proportion of startups producing at least one eligible publication, and the number of highly cited publications per startup. Measures of scientific influence included cumulative citation counts and citation distributions.

Publication practices were assessed through publication type (peer-reviewed publications and preprints), the prevalence and characteristics of collaborative-only publications, and the contribution of preprints to publication output and citation impact. Temporal outcomes included annual trends in startup-led and collaborative-only publication activity.

Additional outcomes examined organizational and geographic variation in scientific publishing activity according to region, founding cohort, valuation, and funding raised. Author-level outcomes characterized affiliation profiles among startup-affiliated researchers, including startup-only, startup-plus-academia, and startup-plus-other affiliations, as well as participation patterns within highly cited publications.

As an external benchmark of institutional scientific critical mass, we assessed whether any AI unicorn startup appeared in the most recent science-wide institutional citation ranking database, which includes only organizations that meet predefined eligibility criteria based on affiliated author-researchers with at least 5 indexed publications (*10*). This comparison was used descriptively to contextualize the scientific scale of AI unicorn startups relative to research-performing organizations.

Finally, outcomes related to the distribution of scientific activity and influence included publication concentration, citation concentration, highly cited paper concentration, authorship concentration among startup-affiliated researchers, and the number of recurrent authors within firms.

### Statistical analysis

All analyses were conducted at the firm level unless otherwise specified. Publication and citation distributions were summarized using counts, proportions, means, medians, interquartile ranges (IQRs), and maxima. Descriptive statistics were reported for the complete startup sample and for publishing startups only. Citation distributions were additionally summarized separately for peer-reviewed publications and preprints.

The primary outcome, the proportion of startups producing at least one highly cited publication, was summarized descriptively for the full sample and across firm characteristics. Secondary outcomes included publication volume, highly cited paper counts, cumulative citations, publication typologies, preprint activity, authorship characteristics, and scientific concentration.

Scientific concentration was quantified using Gini coefficients, Lorenz curves, and top-share statistics for publication volume, cumulative citations, highly cited papers, and author contributions. To distinguish organizational from individual concentration, concentration metrics were calculated at both the startup and author levels.

Temporal analyses examined annual publication activity between 1998 and 2025. Publications were categorized as startup-led (startup-affiliated first author, last author, or alphabetical authorship) or collaborative-only (startup-affiliated authors appearing exclusively in middle-author positions). Startup publication trajectories were compared with the broader AI literature using annual publication counts from Web of Science, INSPEC, and the Preprint Citation Index obtained through a predefined AI bibliometric search strategy (Supplementary Material X). Startup participation was expressed both as absolute publication counts and as a proportion of all AI publications indexed in each year.

Descriptive analyses were conducted according to geographic region, valuation tier, funding raised, publication typology, and author-affiliation profile. Publication typologies were used to distinguish firms according to publication productivity and highly cited output. Valuation tiers were defined as exactly US$1 billion, >US$1 billion to US$5 billion, >US$5 billion to US$10 billion, and >US$10 billion. Funding raised was divided into tertiles.

Given the highly skewed distributions of publication and citation counts, non-parametric statistical methods were used throughout. Mann–Whitney U tests compared two-group distributions, including US versus non-US firms and selected comparisons involving preprint activity. Kruskal–Wallis tests compared publication and citation distributions across valuation tiers, funding tertiles, geographic groups, and founding cohorts. Chi-square tests evaluated associations between categorical publication outcomes and firm characteristics, whereas Fisher’s exact tests were used when expected cell counts were small. Spearman’s rank correlation coefficients quantified monotonic associations between publication activity, highly cited paper production, citation volume, valuation, funding raised, founding year, and firm age.

Author-level analyses characterized affiliation profiles, authorship positions, and participation in highly cited publications. Differences between the full author population and authors contributing to highly cited papers were assessed descriptively. Author concentration was quantified using frequency distributions and Gini coefficients to determine whether highly cited outputs were driven by a small number of recurrent contributors.

As an exploratory extension of the author-concentration analysis, we counted recurrent authors within each firm. Authors linked to multiple startups were assigned to their primary firm, defined as the first startup listed in the startup-affiliation field; a sensitivity check counting these 81 multi-affiliated authors once for each associated firm changed counts slightly but did not alter the ranking of the top two firms. For these firms, recurrent-author counts were compared descriptively with approximate employee counts from publicly available sources. Employee counts were treated as contextual indicators rather than formal analytical variables and were not used to infer employees’ internal scientific or technical contributions beyond the formal literature.

All analyses were conducted programmatically using custom Python scripts and are available at the following repository: https://github.com/q5loisel/unicorn-AI-startup-publications. Dataset construction, affiliation processing, bibliometric analyses, and statistical testing were automated to ensure reproducibility. Monetary values were converted to US dollars using contemporaneous exchange rates. The study data are available at the following repository: https://osf.io/wtnm6.

## Results

### Uneven participation in scientific publishing

The final dataset (see Supplementary Material: Methods and Materials) comprised 317 AI unicorn startups headquartered in 25 countries, collectively associated with 2,077 qualifying scientific outputs, including 1,389 peer-reviewed publications and 688 preprints. Among them, 132 (6.4%) received at least 200 citations and comprise the set considered as highly cited.

Publication activity was highly uneven across firms (Fig. 1A). More than half of startups produced no qualifying output (166 of 317; 52.4%), and most others produced only a small number of papers (median, 4), while a limited set of firms maintained sustained publication activity (maximum, 243).

**Fig. 1.**
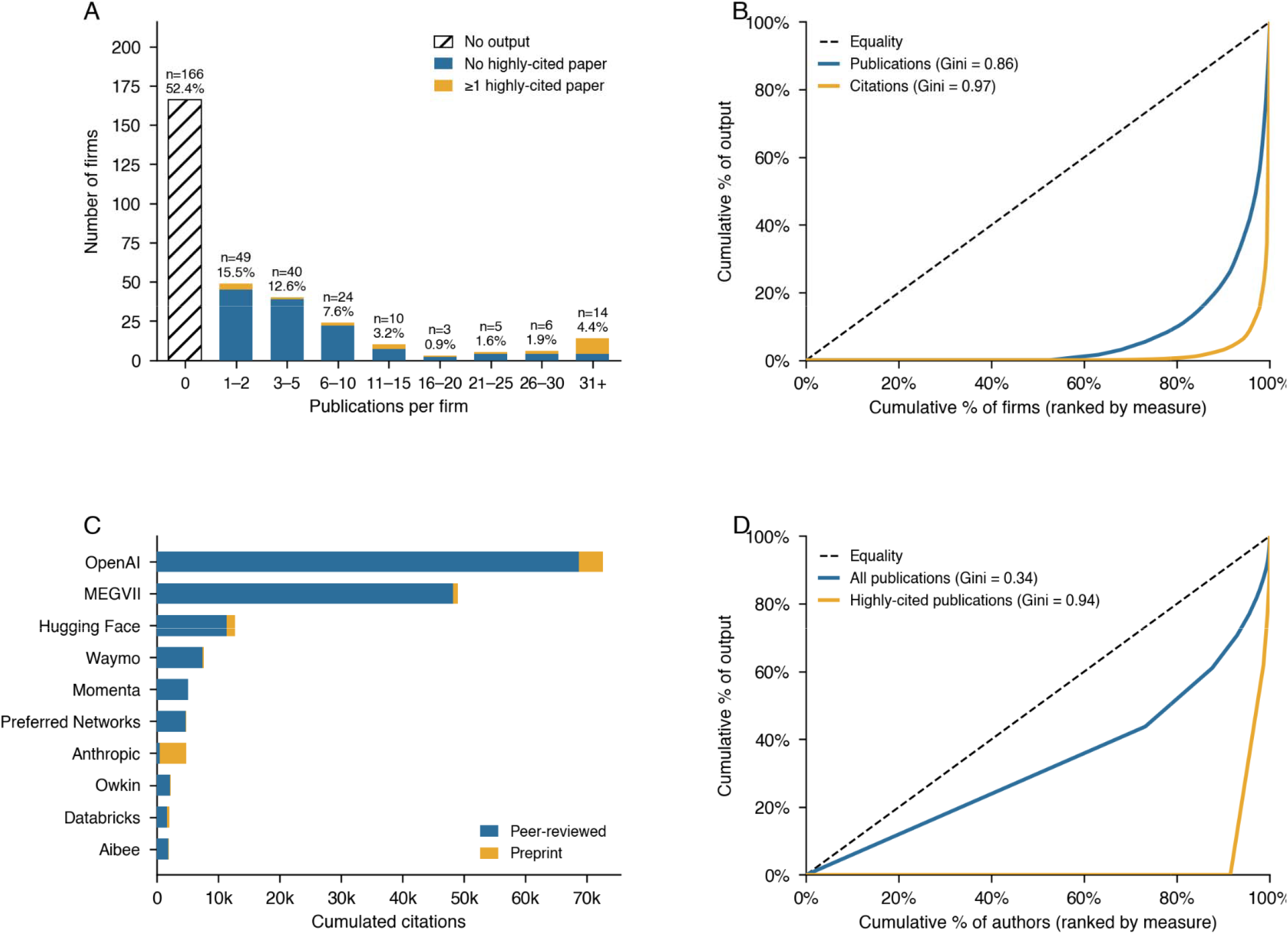
Firm distributions and concentrations of scientific output. (A) Distribution of publications per firm. (B) Lorenz curves: firm concentrations of publications and citations. (C) Top 10 firms by cumulative citations. (D) Lorenz curves: author concentrations of publications and highly cited publications.

This inequality became even more pronounced when examining scientific influence rather than publication volume alone (Fig. 1B). Lorenz curves demonstrated extreme concentration for both publications and citations, with greater inequality for citations (Gini = 0.97) than for publication counts (Gini = 0.86). The bottom half of firms accounted for essentially none of the observed publications or citations. Conversely, the top 5% of firms accounted for 59.9% of all publications and 92.0% of all citations, whereas the top 10% accounted for 76.4% and 96.8%, respectively.

These patterns indicate that published scientific output in the AI startup sector is concentrated among a handful of firms, and that citation impact is even more concentrated than publication activity itself. The degree of concentration observed here exceeds inequalities commonly reported in scientific production systems more broadly (*8, 9*).

Fig. 1C shows the ten most-cited startups, which cumulatively accounted for 88.2% of all citations identified in the dataset. OpenAI alone represented 39.4% of all citations, followed by MEGVII (26.6%) and Hugging Face (6.9%). Highly cited papers were similarly concentrated. Only 24 firms (7.6%) produced at least one highly cited qualifying paper, whereas 293 firms produced none. Among firms with highly cited outputs, the median number of highly cited papers was 1.5, although MEGVII alone accounted for 53 highly cited papers and OpenAI for 30.

Publication strategies varied substantially within the highly cited subset. Anthropic was the clearest top-cited exception, accumulating 90.6% of its citation impact through preprints, whereas most other top-cited firms derived most of their citations from peer-reviewed publications (Fig. 1C). OpenAI, MEGVII, Waymo, Momenta, Preferred Networks, Owkin, and Aibee each received more than 94% of citations from peer-reviewed outputs; Hugging Face and Databricks were more mixed, with 10.7% and 16.6% of citations from preprints, respectively. Preprints had lower average citation counts than peer-reviewed papers but still displayed a heavy-tailed distribution: preprints had a median of 4 citations and a maximum of 1,400, compared with a median of 5 and a maximum of 24,931 for peer-reviewed outputs. Among all publishing firms, the median preprint share was 20.0% of output (IQR: 0–50%), and 24.5% relied on preprints for more than half of their scientific output. These findings suggest that preprints are used unevenly by these unicorns in AI research.

### Authors affiliated with AI unicorns

Across all papers, 1,988 unique startup-affiliated authors were identified: 53.5% held affiliations exclusively within their startup, 38.0% maintained concurrent affiliations with a research institution, and 8.5% co-listed another non-academic affiliation. These groups were represented in 1,204 (58.0%), 1,039 (50.0%), and 230 (11.1%) of the 2,077 papers, respectively. The 132 highly cited papers involved a more narrowly embedded author pool (n = 169): the proportion holding startup-only, startup plus academic, and startup plus other affiliations was 53.3%, 43.2%, and 3.6%, and these groups were represented in 89 (67.4%), 75 (56.8%), and 8 (6.1%) of the highly cited papers, respectively.

Authorship was moderately concentrated across all counted author-publication observations, with 3,324 observations distributed across 1,988 authors (Gini = 0.344). In contrast, highly cited author-publication observations were much more concentrated: although 142 of the 169 authors with at least one highly cited observation appeared only once in this category (84.0%), the distribution across all 1,988 counted authors was highly unequal (Gini = 0.935; Fig. 1D).

The most prolific highly-cited author was associated with 23 highly cited papers, whereas the most prolific author in the full dataset was associated with 48 papers. Twenty-seven repeat HC-paper authors, including 13 with concurrent academic affiliations, accounted for 87 of 229 (38.0%) HC author-publication observations.

None of the startups appeared in a science-wide citation ranking database of organizations and institutions, which requires a minimum critical mass of affiliated author-researchers with at least 5 papers within an organization for inclusion (*10*). Thus, even among the few AI unicorns with measurable citation impact, scientific output appears to be generated by a relatively small set of repeat leading authors rather than being broadly distributed across the workforce. MEGVII and OpenAI had the highest numbers of authors with at least one eligible publication, with 190 and 123 authors, respectively. However, only 8 MEGVII authors and 8 OpenAI authors had five or more eligible publications, corresponding to 4.2% and 6.5% of their author pools. These figures are small relative to the firms’ overall workforces: OpenAI reportedly had approximately 4,500 employees in 2026, while MEGVII has been described as having approximately 2,300 to over 3,000 employees (*11*–*13*). Almost all employees of AI unicorns are decoupled from contributing in leading ways to the scientific literature, despite their credentials, skills, and scientific know-how.

### Scientific output and financial measures

Scientific publishing activity varied substantially across firms with similar financial profiles (Fig. 2A). Valuation was not associated with publication productivity: the Spearman correlation coefficient between firm valuation and total publications was negligible and non-significant across the full sample (ρ = 0.017, p = 0.767, N = 317), including when restricted to publishing firms alone (ρ = 0.100, p = 0.222, n = 151). The association between firm valuation and the number of highly cited papers was similarly weak and non-significant across the full sample (ρ = 0.042, p = 0.457) and remained so also among publishing firms only (ρ = 0.072, p = 0.381).

**Fig. 2.**
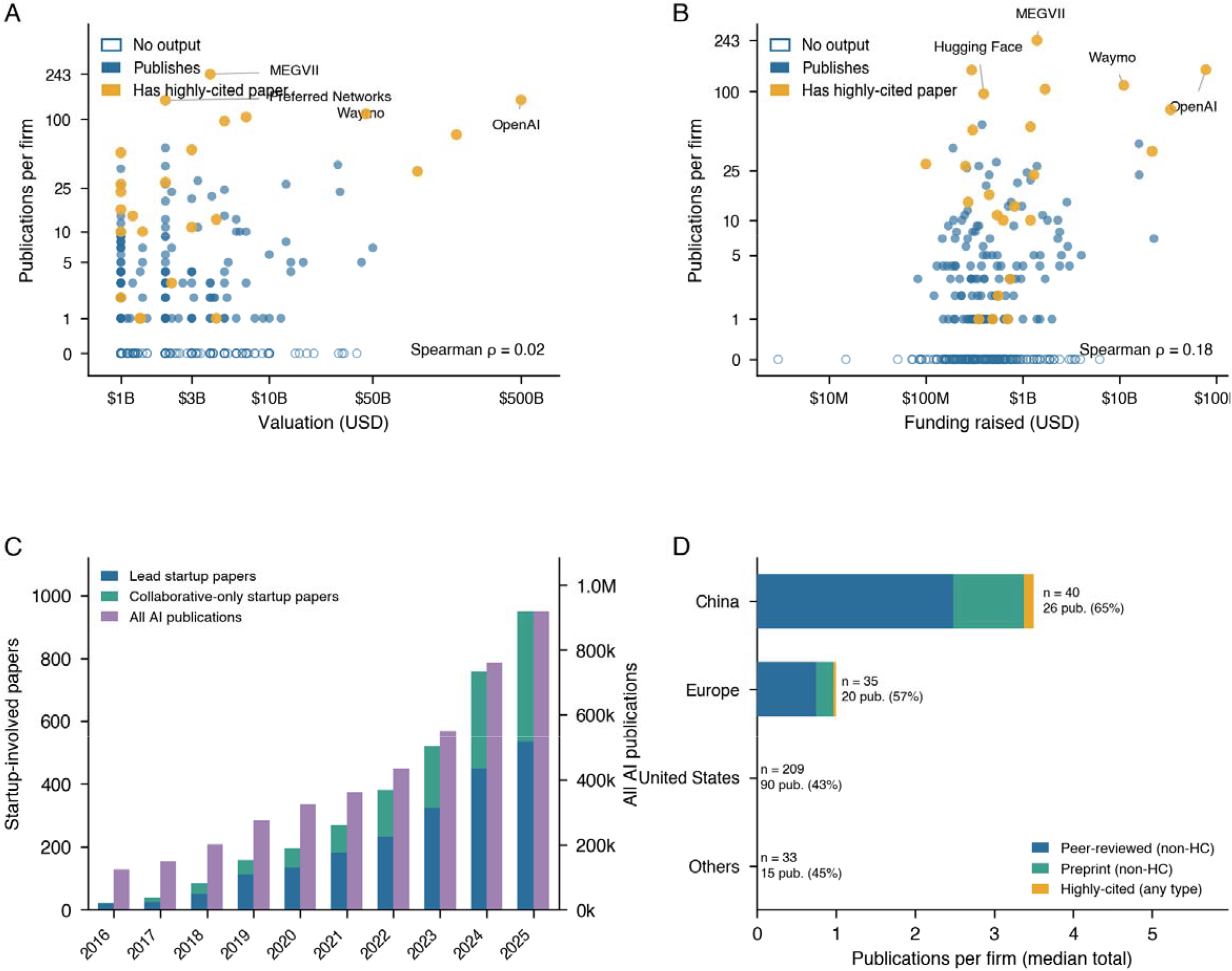
Economic scale, temporality, and geography of scientific activity. (A) Firm valuation versus publication count; (B) Funding raised versus publication count; (C) Annual publication volume evolution for lead and cooperative startup papers, and overall, AI fields (2016–2025). (D) Median publications per firm by geographic region.

Funding raised (Fig. 2B) showed a weak but statistically significant association with publication count (Spearman ρ = 0.181, p = 0.001, n = 315) and number of highly cited papers (Spearman ρ = 0.157, p = 0.005, n = 315), accounting for little of the variation in scientific output. The association was slightly stronger when restricted to publishing firms only, but remained modest for both publication count (ρ = 0.227, p = 0.005, n = 149) and highly cited papers (ρ = 0.190, p = 0.020, n = 149). Although better-funded firms were somewhat more likely to publish and produce highly cited papers, impactful scientific publishing remained concentrated within a small subset of organizations rather than scaling uniformly with financial magnitude.

### Change over time and share in the entire AI literature

Scientific publication involvement among AI unicorns expanded substantially over the study period, but these firms cumulatively remained tiny, negligible contributors to the overall AI published literature (Fig. 2C). Startup-led papers increased from 18 in 2016 to 534 in 2025, while collaborative-only papers, in which startup-affiliated authors appeared in middle-author positions without first, last, or alphabetical authorship, increased from 5 to 416. Overall, AI publications involving startups rose from 23 papers in 2016 to 950 in 2025. These numbers are dwarfed by the broader AI literature, which increased from 123,383 publications in 2016 to 919,486 in 2025. Overall, the share of all AI publications involving these startups remained tiny, ranging from 0.019% in 2016 to 0.103% in 2025.

### Differences across countries

Geographic region was associated with substantial differences in publication output (Fig. 2D). Chinese firms demonstrated the highest median publication count, even if it was still very sparse (3.5 papers per firm); European firms had a median of 1.0 paper, and firms from other regions and the United States had a median of 0 papers (regional differences Kruskal-Wallis H = 13.78, p = 0.003). Chinese firms also had the highest publishing rate, with 26 of 40 firms publishing (65.0%), compared with 20 of 35 European firms (57.1%), 15 of 33 firms in other regions (45.5%), and 90 of 209 US firms (43.1%). These regional differences suggest that organizational context and scientific norms may vary substantially across geographies, with China startups typically publishing some papers, while US startups typically show absolutely no interest in the scientific literature.

## Discussion

Together, these findings suggest that participation in scientific publishing among AI startups is overall very sparse and highly selective and shaped by divergent organizational approaches to scientific visibility and disclosure. A few firms appear to be embedded, albeit to a limited extent, in the traditional scientific communication system, whereas the majority do not engage in formal publishing at all, despite substantial financial scale and technological prominence. Cumulatively, AI unicorns have a negligible presence in the scientific literature. They are the frontrunners in development in the field but show almost total indifference to the usual ways of communicating science.

A particularly notable finding is the absence of a meaningful relationship between company valuation and scientific influence. Firm valuation was not associated with publication productivity or highly cited output, while funding raised showed only weak associations with these outcomes. Again, this shows the near-complete disconnect between traditional ways of disseminating science and what matters in the competitive market environment. One may argue that scientific visibility and impact should contribute substantively to commercial success. If so, investors who try to decide where to invest and valuators could recognize that having peer-reviewed and validated strong science can enhance the chances of successful applications and translations with commercial value. One may also argue that commercial success could also, reciprocally, promote scientific visibility by attracting and enhancing collaborations with highly influential publishing authors and institutions. However, our data suggest an almost perfect lack of any relationship between finances and scientific output. Investors and valuators apparently ignore the peer-reviewed record of these firms and may depend entirely on private information, and other heuristics, or instinct (if not, outright gambling). Powerful firms, on their side, may feel that they can attract talent and skills, bypassing the peer-reviewed publications and citation signaling that energized academics.

This pattern should be considered alongside broader developments in frontier AI, including the consolidation of computing resources, technical talent, and advanced model development within a limited number of firms (*1, 14*–*16*). AI technological advances are apparently expanding fast beyond the boundaries of traditional scientific fora. These patterns are important because scientific publications serve not only to disseminate findings but also to facilitate external scrutiny, the development of cumulative knowledge, and transparency regarding methods, capabilities, and limitations.

Low and uneven participation in the scientific literature has also been documented previously for startups in other sectors (*4*–*6*). Whether frontier AI firms should engage extensively in scientific publishing as a formal societal requirement remains also a matter of debate. Organizations may reasonably choose to protect proprietary innovations and competitive advantages. Nevertheless, in domains where AI systems can have substantial societal consequences, including health-related applications, limited engagement with scientific communication may reduce opportunities for independent evaluation, validation, and cumulative learning. Costly, ineffective, and harmful applications and technologies may become widely used. Even for non-health applications, AI technologies may have other major societal impacts, e.g., high use of energy, use of (increasingly sparse) water resources for data centers, and high impacts on employment, education, and societal norms and behaviors. Furthermore, traditionally, participation in the scientific literature is considered to foster collaboration and accelerate the diffusion of knowledge. However, the rules of the game may be entirely different in this new influential frontier.

Our findings have implications for ongoing discussions concerning transparency, reproducibility, accountability, and governance in AI research (*7, 17, 18*). If increasingly influential AI capabilities emerge within organizations that engage only sparingly and selectively with formal scientific communication, the public scientific record may affect the trajectory and accountability of AI development. Important advances may occur outside traditional scientific channels, potentially complicating efforts to independently evaluate emerging systems, reproduce technical findings, and assess their societal implications.

Several limitations should be considered. Publication retrieval relied on affiliation-based matching and may underestimate outputs associated with undisclosed affiliations, informal collaborations, or internal industrial research that does not appear in the public literature. Concurrent academic and startup affiliations also complicate attribution of scientific activity to industrial versus academic environments. Citation-based indicators are sensitive to disciplinary publication practices, particularly in AI and computer science, where conferences and preprints play important roles in dissemination. Finally, publication activity and citation impact do not directly measure technological capability, commercial success, or societal impact. Accordingly, this study characterizes participation in formal scientific communication rather than innovation more broadly.

Overall, as private for-profit organizations continue to shape the development of frontier AI systems, understanding how industrial actors participate in, or disengage from and largely ignore, the institutions of science may become increasingly important. Issues at stake include research governance, scientific visibility, and the collective production, accumulation, and synthesis of knowledge. Different models of growth in frontier AI would mean that there can be dramatically different (inclusive or restricted) sets of stakeholders who can meaningfully and informatively participate in the ongoing AI revolution.

## Acknowledgments

This study was preregistered on February 26th, 2026, on OSF (https://osf.io/w2pcj). The following OSF project repository includes deviations from preregistration: https://osf.io/wtnm6. During the preparation of this manuscript, the authors used Claude Opus 4.7, 4.8, and ChatGPT 5.5 for code development and manuscript editing. After using this tool, the authors reviewed, verified, and edited the content and took full responsibility for the final publication.

## Author contributions

Conceptualization: QEAL, ASL, JPAI

Data Curation: QEAL

Methodology: QEAL, ASL, JPAI

Investigation: QEAL

Formal Analysis: QEAL, JPAI

Visualization: QEAL, JPAI

Writing – original draft: QEAL, JPAI

Writing – review & editing: QEAL, JPAI

## Data, code, and materials availability

All data, code, and materials used in the analysis can be found at https://osf.io/wtnm6

